# DigestR an open-source software tool for visualizing LC-MS proteomics data resulting from natural protein catabolism

**DOI:** 10.1101/2024.10.08.617287

**Authors:** Dimitri Desmonts de Lamache, Raied Aburashed, Soren Wacker, Ian A. Lewis

## Abstract

Protein catabolism is an essential biological function supported by every living organism. Although liquid chromatography mass spectrometry proteomics has advanced considerably over the past decade, protein catabolism in natural systems is still difficult to study. One reason for this is the lack of software tools designed specifically for decoding the complex mixtures of peptides that result from *in vivo* protein digestion. To address this, we developed DigestR, an open-source software tool designed specifically for the analysis of LC-MS proteomics data. DigestR allows users to visualize naturally occurring peptides and align them to a reference proteome at display them at either a proteome-wide and protein-specific level. These visualization tools allow users to track the patters of peptides occurring in natural systems and map naturally-occurring proteolytic cut sites. To demonstrate these functions, we used DigestR to analyze a mixture of peptides resulting from the *in vitro* digestion of human hemoglobin and bovine albumin with a cocktail of well characterized proteases. As expected, DigestR correctly identified both the proteins involved and the proteolytic cut sites produced by our protease cocktail. We then used DigestR to analyze the complex semi-ordered hemoglobin digestion pathway used by the malaria parasite *Plasmodium falciparum.* We show that DigestR successfully identified the proteolytic cut sites linked to the Plasmepsins, a protease known to be involved in hemoglobin digestion by the parasite. Collectively, these findings show that DigestR can be used to help visualize and interpret the complex mixtures of peptides occurring through *in vivo* protein catabolism. DigestR can be downloaded from www.lewisresearchgroup.org/software.

## Introduction

Protein catabolism, the breakdown of proteins into their constituent peptides or amino acids, is an essential biological function that directly affects a wide range of cellular activities. These activities include supporting metabolic needs of the cell, balancing steady-state levels of proteins to maintain homeostasis, allowing cells to respond to environmental stimuli, and enabling progression through the cell cycle [1–4]. To support these functions, cells use a diverse range of endoproteases (e.g. the proteasome and trypsin), exoproteases (e.g. carboxypeptidases and aminopeptidases) to degrade proteins. These enzymes can act independently of one another and thereby create complex semi-ordered protein digestion pathways, such as the pathway used to degrade hemoglobin in the malaria parasite, *Plasmodium falciparum* [5–9] .

The analytical tools needed to study complex naturally occurring proteolytic cascades are well established. Modern liquid chromatography mass spectrometry (LC-MS) can detect tens of thousands of peptides [10, 11] and has limits of detection approaching the attomole level under idealized conditions [12, 13]. However, these tools have largely been designed to address the needs of traditional proteomics studies, which use a workflow wherein proteins are digested *in vitro* using one of a few known proteases (lysC, trypsin, etc.). This workflow greatly simplifies downstream informatics as it restricts the search space to a handful of expected proteolytic fragments. However, the semi-ordered proteolytic cascades occurring in natural systems produce large collections of closely related peptides that do not have well defined cut sites that can be defined *a priory* [14]. As a result, novel “digestomics’ informatics strategies must be employed to track these proteolytic cascades [15].

Although there are dozens of powerful proteomics software tools available (e.g. SeaQuest, Mascot, RawTools, MaxQuant, ProteomeDiscover, etc.) [16–19], there are limited tools available that meet the unique challenges of analyzing complex mixtures of naturally occurring peptides. Currently, the best available tools in this space include Peptigram [20], iPiG [21], PeptideExtractor (PepEx) [22] and EnzymePredictor [23]. Though powerful for *in vitro* protein digestion analyses, these software tools rely on a defined precursor-product relationship to function and are thereby not well suited to analyzing semi-ordered proteolytic cascades occurring *in vivo*.

The key features needed to support the analysis of these proteolytic cascades include: 1) an ability to handle peptides of any length and with any carboxy or n-terminal cleavage, 2) a mechanism to align peptides with a reference proteome and visualize the naturally-occurring peptides in a genomics context, and 3) a scoring system for differentiating real-versus spurious matching (an problem inherent to proteome-wide alignment when searches are computed with minimal constrains) [15]. To address these challenges, we developed a new open-source graphics-based software tool (DigestR) that is specifically designed for untargeted peptide mapping and interpreting complex proteolytic cascades that occur from the natural digestion of proteins. We designed DigestR to enable peptide visualization in a genomic context, provide visual readouts of peptide accumulation, and provide a variety of data overlay and analysis tools to help support the interpretation of proteolysis. DigestR was built in the R statistical software environment (freely available from www.r-project.org), and was adapted from the open source rNMR data analysis package for metabolomics [24]. DigestR is freely available from www.lewisresearchgroup.org/software. We illustrate the function of this tool using both well-defined *in vitro* digestion experiments as well as the analysis of the semi-ordered *P. falciparum* hemoglobin digestion cascade.

## Experimental section

### *In vitro* sample preparation

To test our software, we digested purified Human hemoglobin (Sigma, H7379) as well as Bovine serum albumin (BSA; Sigma) with different proteases used either individually or in combination. Trypsin, chymotrypsin and GluC were acquired from Thermo-Fisher scientific (PI90057, PI90056 and 90054 respectively). Briefly, proteins were resuspended in digestion buffers (**supplemental Table 1**) to a concentration of 0.5 mg/mL. Proteases were then added to a final protease: protein ratio of 1:20 (w/w) and incubated for seven hours at the optimal temperature for each enzyme (**supplemental Table 1**). Enzymatic reactions were stopped by adding formic acid to a final concentration of 1%. Samples were subsequently evaporated using an Eppendorf 5301 Vacufuge Concentrator System (Marshall Scientific, EVCC) and stored at -80℃ until mass spectrometry analysis.

### *P. falciparum* strain and sample preparation

The use of human blood for this work was approved under the Conjoint Health Research Ethics Board (REB21-1835). Wildtype *P. falciparum* (3D7) parasites were cultured at 2% hematocrit and <5% parasitemia in human blood incubated in RPMI (Fisher Scientific, cat. no. 31-800-022) supplemented with 25 mM HEPES (pH 7.4; VWR, cat. no. 97061-822), 0.1 mM hypoxanthine solution in 1M NaOH (VWR, cat. no. A11481-14), 100 µM gentamycin (Gibco, cat. no. 15750060), 600 µM sodium bicarbonate (Millipore Sigma, cat. no. S5761-1KG), and 0.25% AlbuMAX™ II Lipid-Rich BSA (Gibco, cat. no. 11021037). Parasites were incubated at 37°C with 5% CO_2_ and 5% O_2_ in a humidified tri-gas incubator. Prior to the experiment, parasites were synchronized twice with a 5% sorbitol solution, grown to 5-10% parasitemia, and allowed to reach the trophozoite stage. Trophozoites were purified using a 30%/70% Percoll gradient (centrifugation at 3500 xg for 15 minutes) and incubated with uninfected red blood cells at a 5:1 (uninfected: infected cells) ratio for 12 hours to allow for reinvasion. Following reinvasion, parasites were washed with PBS three times (5 minutes at 2500 RPM) and resuspended at a hematocrit of 0.2%. Peptides were extracted at 36 hours post-invasion in an ice-cold 90% methanol solution, centrifuged at 10,000 RPM at 4℃ for 10 minutes. Supernatants were transferred to a clean Eppendorf tube for storage at -20℃ until proteomics analysis.

### LC-MS Analysis

All LC-MS data were acquired at the Calgary Metabolomics Research Facility (CMRF). Proteomics samples were evaporated in a speedvac and resuspended in 100 µL of a resuspension solution (5% HPLC-grade acetonitrile 0.1% formic acid in LC-MS-grade water). Subsequently, samples were vortexed and sonicated. Protein concentration was determined with a Nanodrop (280 nm absorbance) and adjusted to 10 µg/mL. Samples were C18-cleaned up and eluted in 1 mL of elution solution (80% HPLC-grade acetonitrile and 0.1% formic acid in LC-MS-grade water). Finally, samples were dried and resuspended in a reconstitution buffer (5% acetonitrile, 0.1% formic acid in HPLC-grade water) to a final concentration of 1 µg/µL. Samples were analyzed in positive mode on an Orbitrap Fusion™ Lumos™ Tribrid™ Mass spectrometer. Samples were eluded using nanoflow 300 nl/min using a 120-minute reverse phase (C18) gradient. The source voltage was set to 2.5 kV and the transfer tube was set to 275 °C. MS1 scans were acquired in the Orbitrap at a 120,000 resolving power with a range of 350-1600 m/z, an S-Lens RF of 30%. The top 15 peptides in each MS1 scan were selected for data dependent MS/MS fragmentation by CID at 35% NCE and were detected on the ion trap at a resolving power of 15,000.

### Data processing

Thermo Fisher LC-MS/MS raw data were converted using ProteoWizard 3.0.18312 MSConvert (**see Table 1 for settings**). Peptides were then identified using Mascot Daemon using the settings outlined in **Table 2**. For in vitro experiments, all searches were performed against either a full *H. sapiens* proteome or a full *B. taurus* proteome in the Mascot server. One important difference between established proteomics processing and the DigestR processing method is that the former usually filters out peptides shorter than six amino acids. However, naturally occurring peptides can be shorter and our searches parameters allowed peptides >2 in length to be analyzed. Following peptide identification searches, files were exported as .csv and then imported into DigestR. Files can be processed individually or in batch, allowing time intensive operations to be performed with minimal user interaction. DigestR files are saved in .dcf format which can be opened, viewed, and analyzed at the user’s convenience.

**Table 1.** Mascot Daemon parameter settings. Mascot Daemon user search parameters used for peptide scoring and identification.

| Setting | Value |
| --- | --- |
| Enzyme | None |
| Max. missed cleavages | 2 |
| Peptide charges | 1+, 2+, 3+ |
| Peptide tolerance ± | 20 ppm |
| MS/MS tolerance ± | 0.6 Da |
| MS/MS ion search | On |
| Variable modification | Deaminated (NQ)<br>Oxidation (HW)<br>Oxidation (M)<br>Trioxidation (C) |
| Data format | Mascot generic |

**Table 2.**
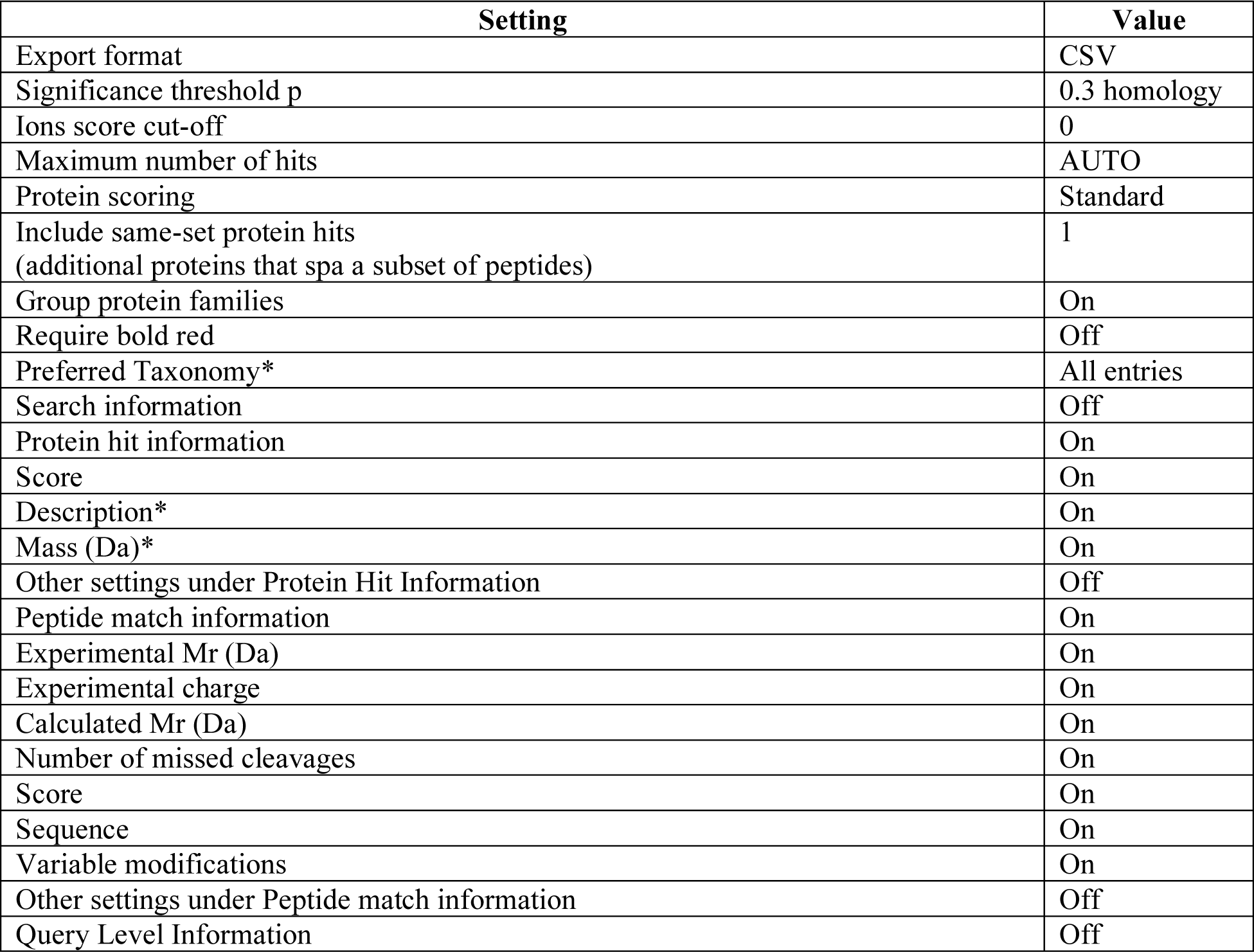
Mascot Daemon export search result settings for DigestR analysis. Settings used to export search results from Mascot Daemon. This file format enables data to be imported into DigestR

## Results and discussion

### Digestion mapping

One of the primary complications in analyzing peptides from naturally occurring proteolytic cascades is that peptides can arise from the action of many different endo- and exoproteases that produce a complex ensemble of related peptides. These ensemble phenotypes represent the total system of the entire proteolytic system, rather than the action of a single enzyme. To visualize this, DigestR uses a coincidence scoring system wherein all peptides, regardless of their relative abundance within the digest, are aligned to the entire proteome at an amino acid level. Coincidence scoring for digestomics is described in detail elsewhere [15]. Briefly, digestion mapping using relatively permissive peptide match thresholds and allows short peptides (>3 residues) to be included in the search. Though essential for understanding protein digestion, these settings result in significant false discovery wherein peptides are spuriously matched across the proteome. To address this, DigestR uses a coincidence scoring strategy wherein peptide matches are expressed as a per-residue average for each protein (Fig 1). Under this algorithm, correctly identified peptides map linearly to the same loci whereas spurious peptide assignments are randomly distributed across the proteome. Signal analysis techniques and reverse/randomized proteome search can then be used to establish signal to noise ratios that define the coincidence score analysis thresholds [15]. The vector of coincidence scores reporting on each protein across the proteome represents digestion phenotype for each sample (Fig 1A). When viewing this digestion map at the proteome level, proteins above the coincidence threshold are displayed automatically (Fig 1A). This automatically determined threshold can be adjusted manually. Peptides that contribute to each protein’s coincidence score can be viewed by toggling between the proteome wide and protein-specific DigestR views (Fig 1B and C).

**Figure 1:**
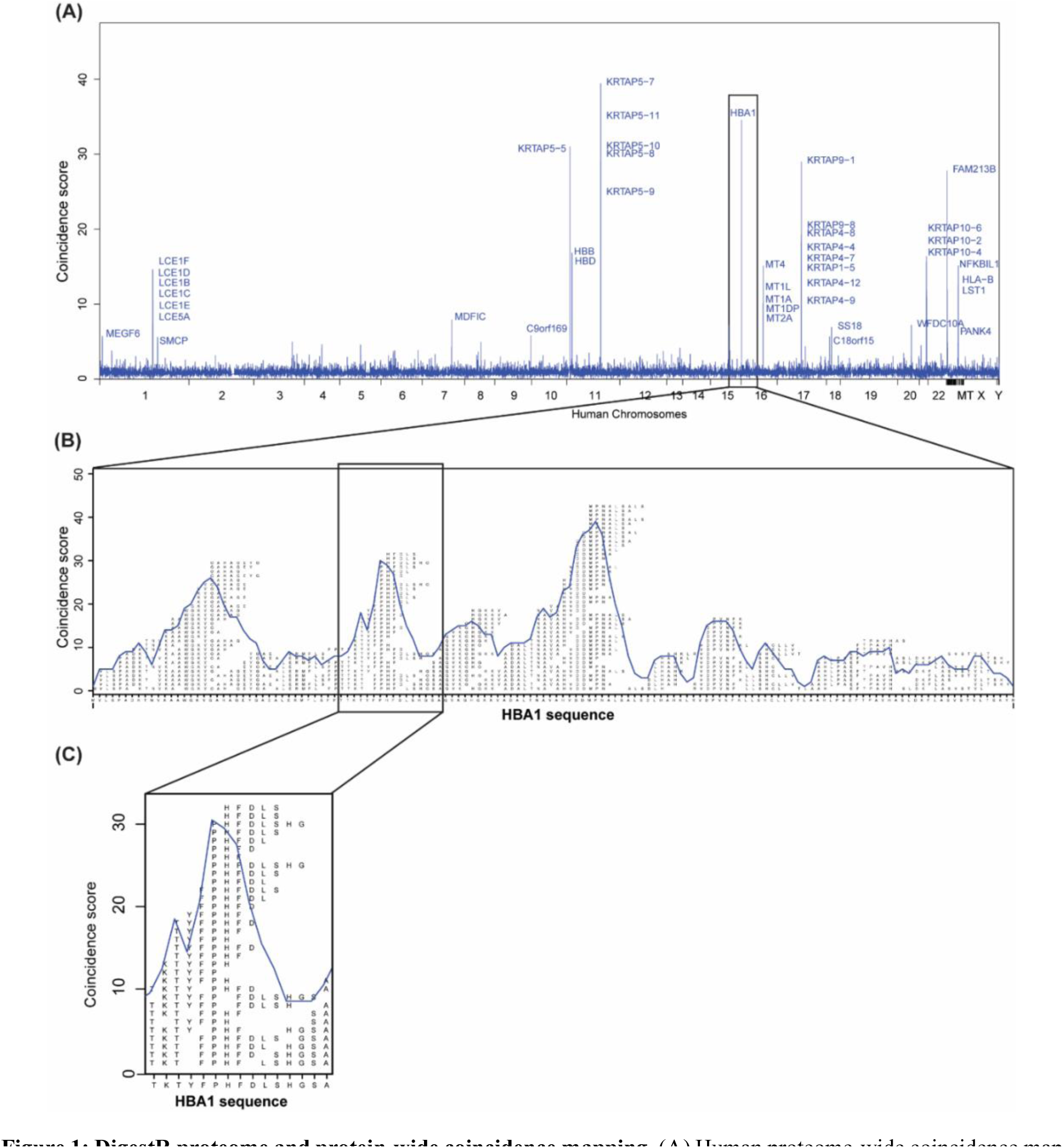
DigestR proteome and protein-wide coincidence mapping. (A) Human proteome-wide coincidence map showing proteins digested by *Plasmodium falciparum* throughout its intraerythrocytic cycle. Coincidence scores are calculated by aligning all potential assignments—regardless of peptide length and including all peptides with Mascot scores of 10 or higher—to their corresponding proteomic loci. The number of peptides aligned to each locus is then represented as an average for each protein. While correctly identified peptides consistently align with the same loci, false hits resulting from incorrect peptide assignments are randomly distributed across the proteome. As a result, averaging the total number of peptides mapped to each locus on a protein-by-protein basis enhances the signal-to-noise ratio. (B) and (C) Protein-wide coincidence maps of human α-hemoglobin digested by *Plasmodium falciparum*.

### DigestR user interface

DigestR GUI was developed to allow users to iteratively interact with datasets. To facilitate data analysis, we created eight graphical windows that allow users to adjust their display settings and view summary data of peptides (Fig 2). These tools include a proteome generation window allowing users to compile proteomics data from BiomaRt to generate a .csv proteome file that will serve as query for peptide alignments (Fig 2A). Following proteome generation, users can import their .csv file generated from mascot and select a proteome of interest to align identified peptides against in the process mascot file window (Fig 2B). In the data visualization window, users can toggle between the proteome view and the protein specific view (Fig 2C and D). Users can also overlay known protease cut sites that they can import from their own .csv file (Fig 2E) and can also overlay and customize digestion map through a data overlay tool (Fig 2F). Finally, DigestR also allows users to perform analyses of peptides size distributions within a digest (Fig 2G), as well as identifying peptide N- and C-termini ends (Fig 2H). These tools aim to facilitate data analysis, help identify unknown cut sites, and anchor the digestion profiles in a biological context.

**Figure 2:**
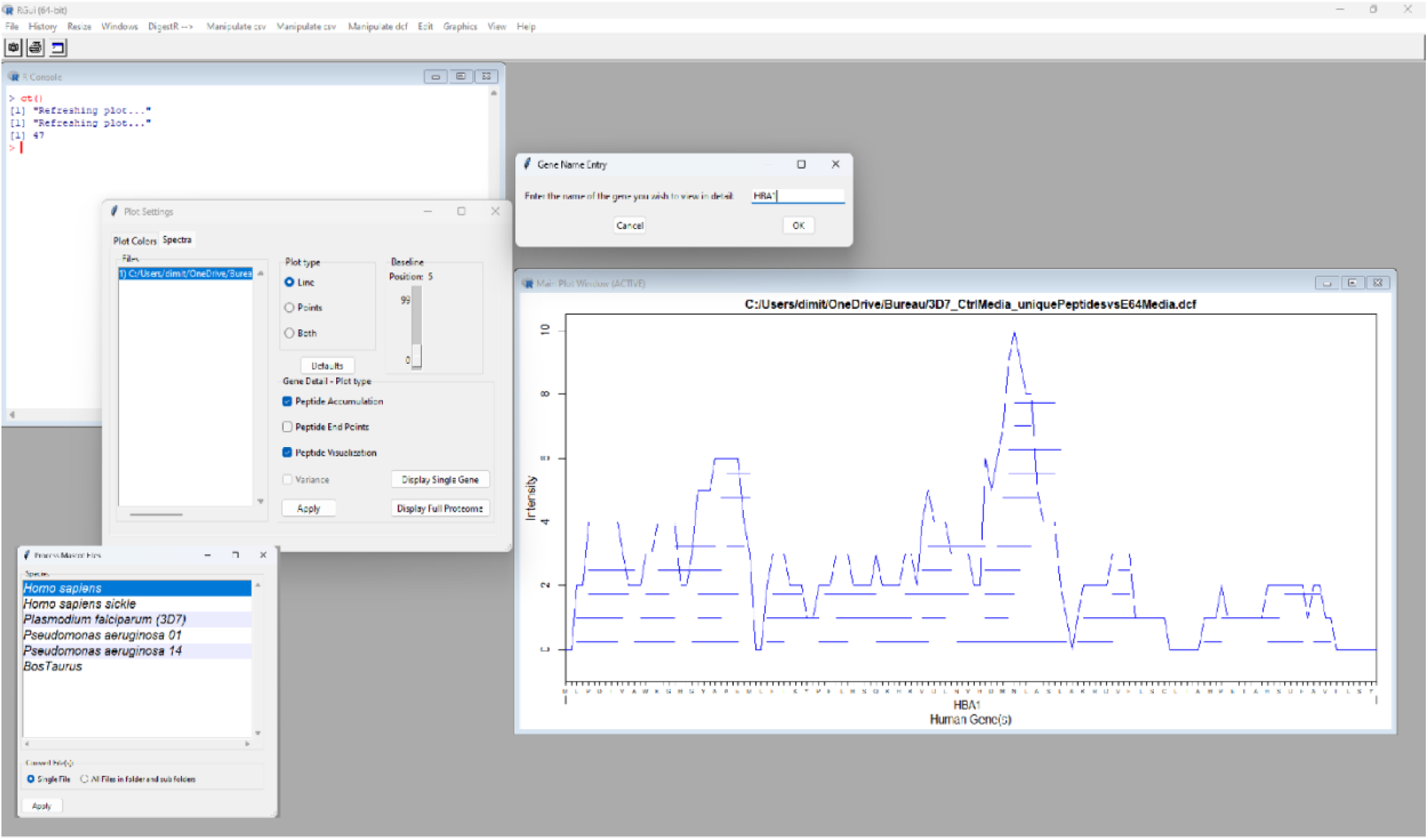
DigestR main plotting window screenshot illustrating several basic features of the program: Users upload .csv files and (a) select a proteome to align peptides against. (b) DigestR performs peptide alignments and generate one-dimensional digestion maps. (c) Proteolytic maps can be displayed either at the proteome wide or protein level by calling for a specific protein. (d) Maps can be overlayed and customized to the user’s needs.

### *In vitro* demonstration study

To assess the performance of digestR, we conducted a series of *in vitro* digestion experiments using *H. sapiens* hemoglobin, and *B. taurus* bovine serum albumin (BSA) and monitored the peptide accumulation patterns produced though *in vitro* digestion. As expected from *in vitro* digestion of these proteins, the proteome-wide views show peptides accumulating at their respective proteomic locations (Fig 3). Coincidence peaks were observed in chromosome 10 and chromosome 16 in the *H. sapiens* proteome, the locations for hemoglobin-α and hemoglobin-β. Similarly, DigestR analysis of *in vitro* proteolysis of BSA showed a coincidence peak at chromosome 6 in the *B. taurus* proteome, the known location for albumin (Fig 3 A and B). To assess the peptides that were contributing to the coincidence peaks, DigestR was toggled to the protein-level view (Fig 4 A and B). Protein-level views showed a proteolytic cleavage pattern characterized by peaks and valleys in the coincidence score resulting from the differential accumulation of peptides at different loci within the protein. Analysis of technical replicates (N=3) showed that the pattern of coincidence peaks was reproducible sample-to-sample (Fig 4C). To determine if digestion maps could be informative about protease activity, we overlayed putative protease cut sites of known enzymes on their respective digestion maps. As shown in **Figure 5**, putative cut sites largely map with corresponding coincidence valleys. This is well illustrated with the trypsin/lysC protease cocktail which exhibits a very strong substrate specificity for a lysine and arginine (Fig 5A). These data demonstrate that peaks observed in the coincidence score report on regions of peptide accumulations whereas valleys represent regions that are subject to proteolytic cleavage.

**Figure 3:**
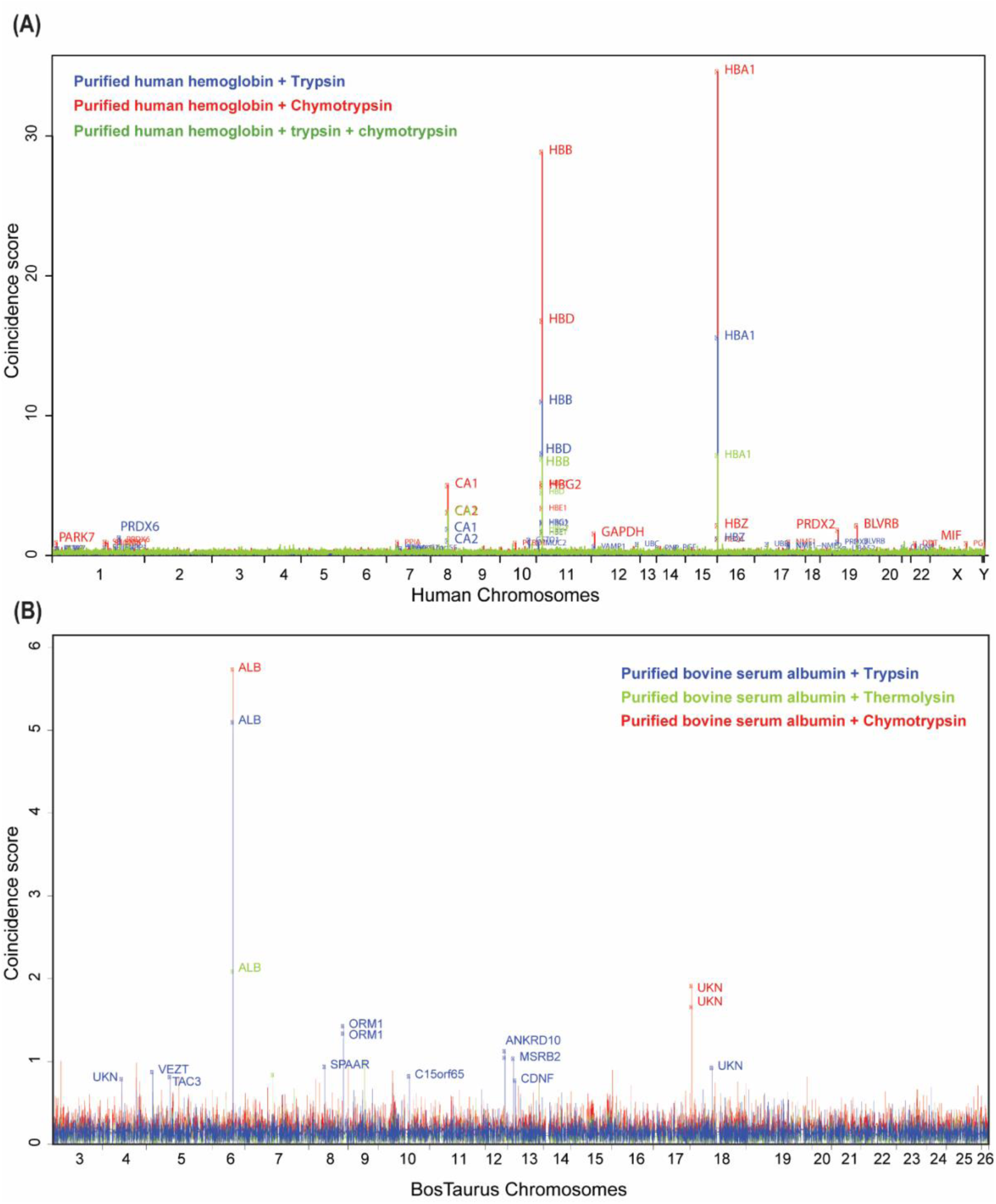
Example of proteome-wide digestion maps. *In vitro* digestion maps of (A) human hemoglobin and (B) bovine serum albumin digested by trypsin (dark blue line), chymotrypsin (red line) and trypsin and chymotrypsin (green line).

**Figure 4:**
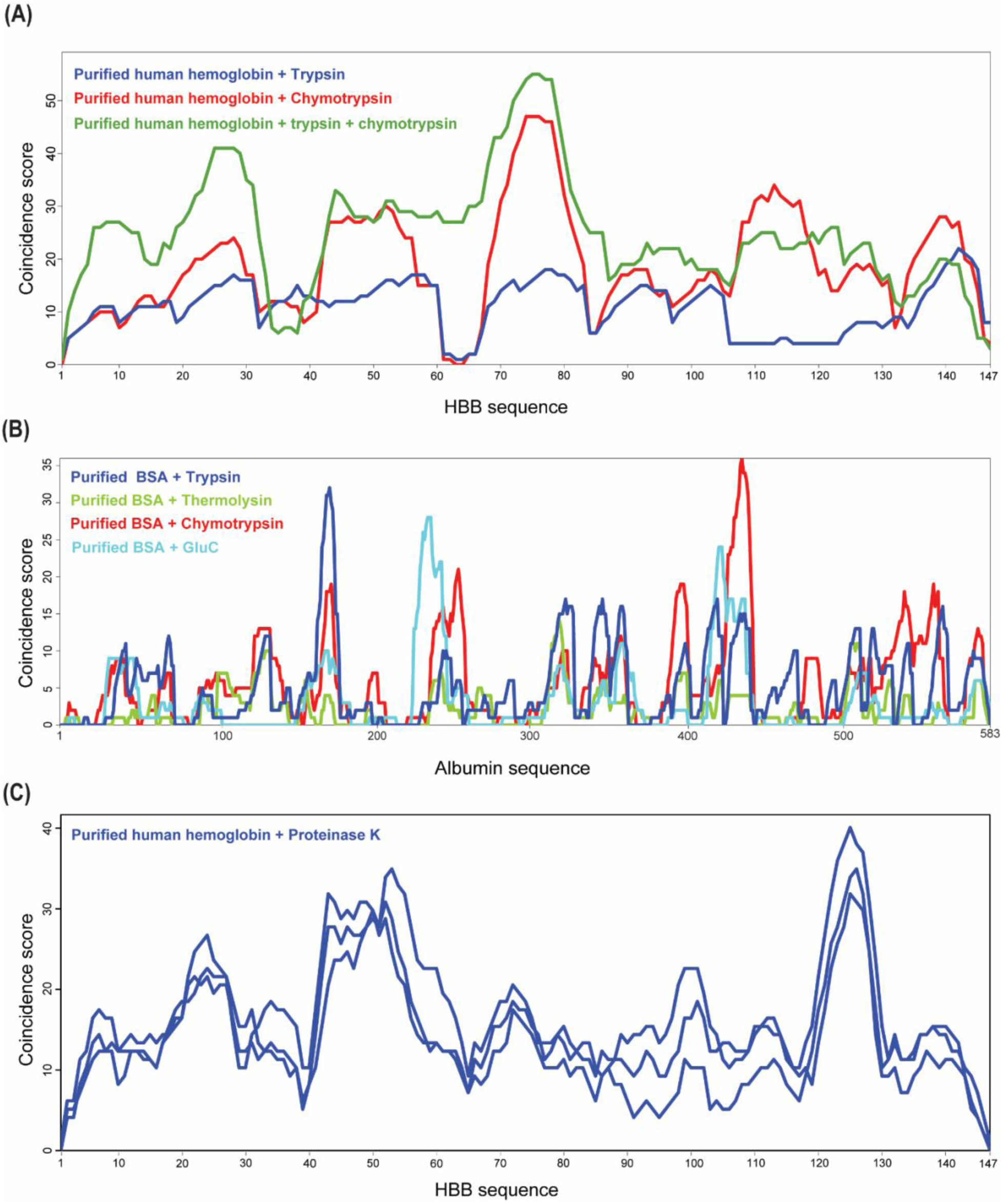
Example of protein level digestion maps. *In vitro* digestion of (A) human hemoglobin and (B) bovine serum albumin digested by trypsin (dark blue line), chymotrypsin (red line) and trypsin and chymotrypsin (green line) and GluC (light blue line). (C) Protein level digestion maps overlay of human hemoglobin digested by proteinase K (n = 3).

**Figure 5:**
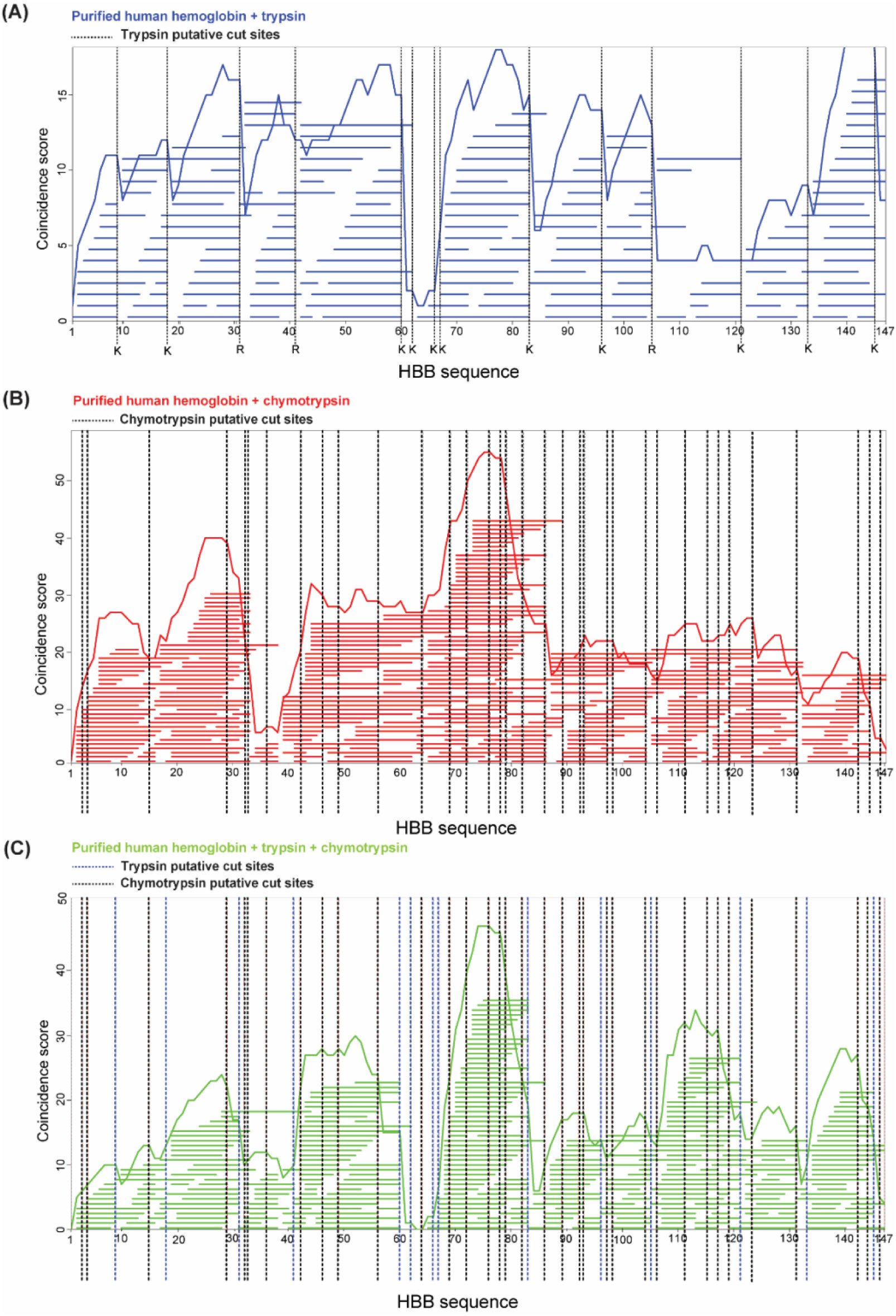
Example of protease putative cut sites overlay on coincidence maps. (A) Digestion map of human β hemoglobin chain (dark blue line) overlayed with trypsin putative cut sites (dashed black lines). (B) Digestion map of human β hemoglobin chain (red line) overlayed with chymotrypsin putative cut sites (dashed black lines). (C) Digestion map of human β hemoglobin chain (gree line) overlayed with trypsin (dashed blue lines) and chymotrypsin putative cut sites (dashed black lines).

DigestR allows users to analyze peptide size distribution as well as their carboxy and amino termini (Fig 6). As expected, carboxy termini of peptides digested with the trypsin/lysC cocktail were heavily skewed towards lysine and arginine residues (46% and 17 % respectively; Fig 6 B). Chymotrypsin-digested peptides were predominantly cut C-terminally at leucine (27%) and phenylalanine (15%), while carboxy termini residues peptides exposed to GluC were dominated by glutamic acid (25.5%) and glycine residues (17.5%; Fig 6 B). DigestR also allows these data to be normalized according to amino acids abundance in the protein sequence. This feature prevents highly represented residues within a protein sequence from skewing the relative percentage of peptide cleavage sites identified within the digests (Fig 6C). Finally, digestR allows for the analyses of protease cut site specificities for the p4-p4’ positions by generating logoplots (Fig 6D).

**Figure 6:**
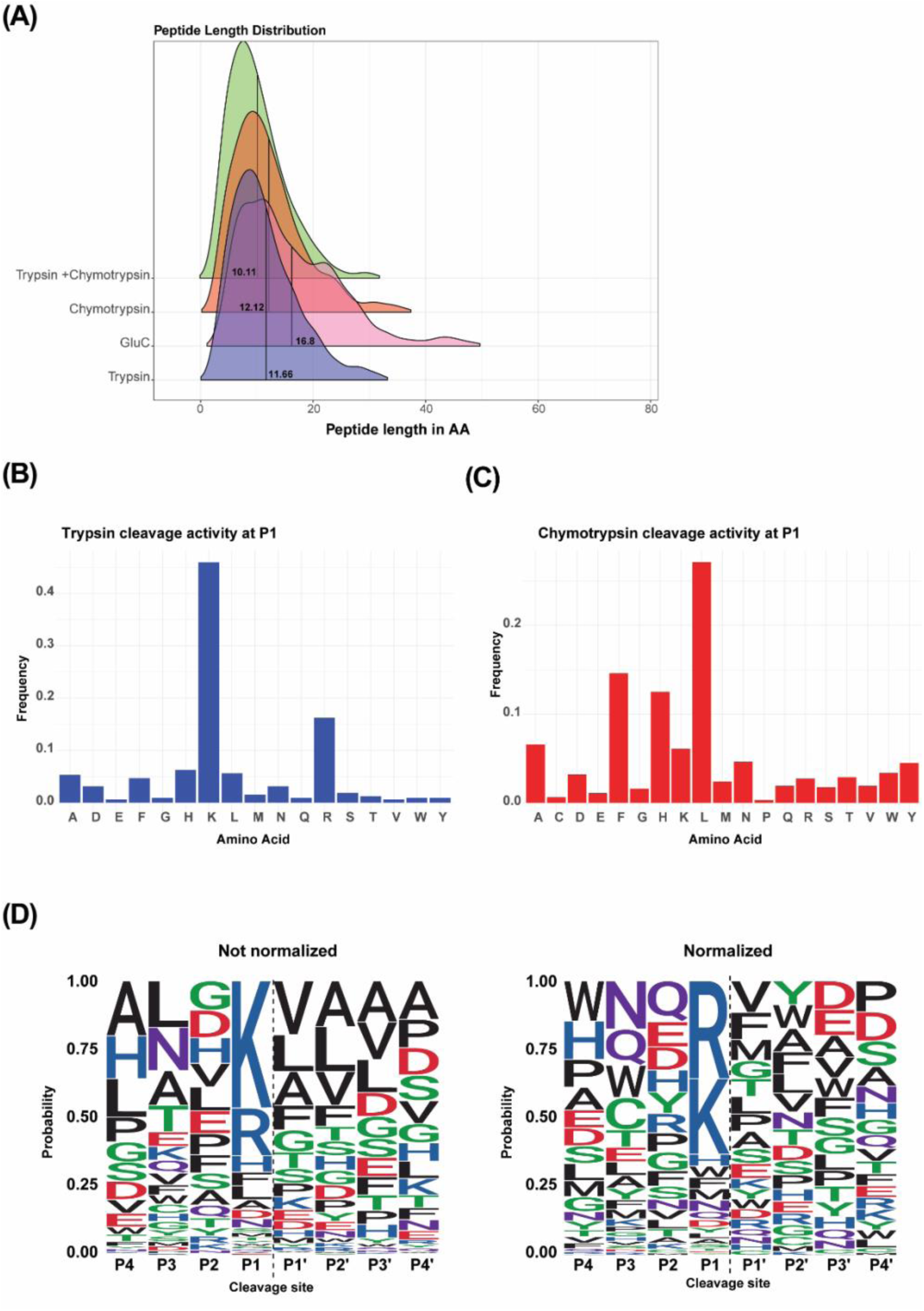
DigestR allows for the analysis of digested peptides. Density plot showing the hemoglobin peptide length distribution (in amino acids) following trypsin (blue), GluC (pink), chymotrypsin (green) (B, C) Amino acid frequency at P1 position following trypsin (B) and chymotrypsin (C) digestion. (D) LogoPlot for the p4-p4’ window following trypsin digestion. The height of the amino acid represents its relative abundance within the peptidome. Amino acid frequencies can be normalized to the protein sequence (left panel).

### Hemoglobin catabolism by *P. falciparum*

To demonstrate that our software could facilitate the analysis of natural protein catabolism *in vivo*, we mapped peptides resulting from the natural digestion of hemoglobin by the *P. falciparum* strain 3D7. Hemoglobin is the most abundant erythrocytic protein making up to 95% of its dry mass [25]. Previous studies have found that *P. falciparum* digests up to 80% of the intraerythrocytic hemoglobin throughout its asexual reproduction cycle [26]. Consequently, most peptides originating from *P. falciparum* digestome should derivate from hemoglobin, which makes the malaria parasite the ideal model system to test our software in more complex biological settings. As expected, digestR proteome wide views showed that most peptides resulting for *Plasmodium*-driven proteolysis during its asexual reproduction cycle were derived from hemoglobin (Fig 7A). Protein views of hemoglobin α and β chains revealed that naturally occurring peptides accumulated around five major loci on the α Hb chain and six loci on β Hb chain (Fig 7B). Examination of the secondary structure of Hb revealed no correlation between cleavage sites and specific secondary structures (Fig 7B). Moreover, our analysis showed that in some cases the valleys in these coincidence maps aligned with published cut sites from *in vitro* analyses of *Plasmodium* proteases [7, 27]. Our data also show that in addition to major cut sites, the pattern of peptides reflected the progressive actions of multiple aminopeptidases resulting in complex “ladders” of closely-related sequences that have been reported previously (Fig 7B) [28]. Investigations of carboxy ends of peptides revealed that *P. falciparum* preferentially cleaved hemoglobin at alanine, leucine and valine residues (Fig 7C). Finally, peptide analyses revealed that hemoglobin peptide length following *P. falciparum* proteolysis averaged 9.3 amino acids in length (Fig 7D). This complex pattern is consistent with the malaria litterature [5, 29, 30].

**Figure 7:**
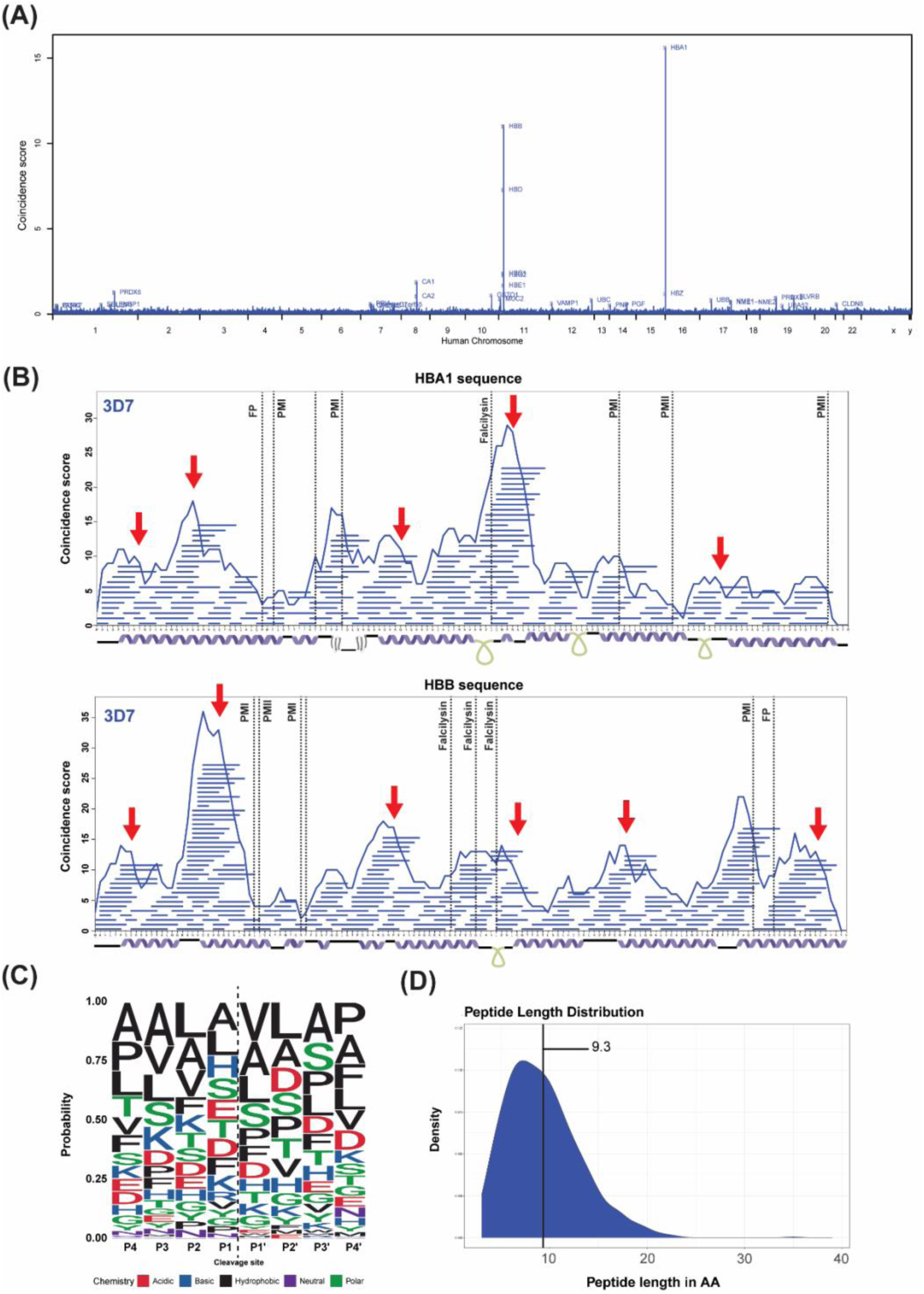
Natural hemoglobin digestion by *P. falciparum*. (A) Proteome view of peptides resulting from natural protein degradation during P. falciparum intra-erythrocytic cycle. (B) Protein level proteolytic maps of hemoglobin α (upper panel) and hemoglobin β (lower panel) digested by *P. falciparum*. Dashed lined indicates known *P. falciparum* protease cut sites. Arrows indicates new putative cleavage sites. (C) Logoplot for the p4-p4’ window for peptides resulting from natural *P. falciparum* cleavage activity. (D) Hemoglobin peptide length distribution following *P. falciparum* proteolysis.

### Challenged solved by digestR and applications

DigestR provides a convenient analysis platform for studying protein catabolism. The peaks and valleys in the coincidence scores provide a simple readout for visualizing the complex biological digestion phenotypes that results from the ensemble action of many proteases. These maps, along with the tools for analyzing peptide numbers, length and distribution, and C- and N-termini distributions provide a toolkit for understanding proteolytic phenotypes. One obvious application of this tool is to help map the proteolytic products that are linked to a specific enzyme. We anticipate that this tool would be a powerful tool asset in the analysis of protease knockouts or protease inhibitors. Herein we illustrate the application of peptide mapping in the context of malaria. However, this host-pathogen dynamic is not singular. Proteolytic mechanisms are known to play a central role in many biological processes through the expression or upregulation of proteolytic enzymes [31–39]. Some specific examples include mucin degradation by the Pic protease, which promotes colonic colonization by *Escherichia coli* and *Shigella flexneri* [31, 32], protein degradation in cystic fibrosis lung sputum by *Pseudomonas aeruginosa* [34, 35], and degradation of the extracellular matrix by cancerous cells for growth, invasion and metastasis [37, 38]. Moreover, it has been found that pancreatic ductal adenocarcinoma cells, access extracellular pools of albumin through macropinocytosis and can use this protein as their sole source of essential amino acids [36]. Peptidomics analysis of plasma from breast cancer patients showed a greater than 4000-fold enrichment in certain peptides [33]. DigestR could also have broader industrial applications, for example in the analysis of fermentation products, characterization of novel enzyme activity, and to understand the digestion of food in livestock. In each of these diverse examples, DigestR will facilitate the research by allowing researchers to visualize and understand complex digestion phenotypes.

### Limitations

Although our analysis demonstrated that the tool faithfully recreates the expected cut sites from proteolytic fragments, users should be aware of some limitations that are inherent to this style of analysis. Firstly, DigestR is qualitative: the heights of the peaks do not correspond to changes in peptide concentration – they report the number of peptides mapped to each locus independent of their abundance. In addition, the valleys in the coincidence scores can have multiple interpretations: they can arise from either the absence of cut sites or the presence of many consecutive cut sites that generate peptides that are too small to be captured by the mass spectrometer. Thirdly, not all peptides ionize evenly, which may produce uneven coincidence scores resulting from the peptide chemistry.

## Conclusion

Despite the important role that protein catabolism plays in the metabolism of organisms, there are currently no publicly available tools designed specifically for characterizing naturally occurring protein catabolism. Herein, we demonstrate that DigestR’s graphics toolkit can help map proteolytic cut sites and decipher complex proteolytic phenotypes. DigestR’s alignment tools provide a window into the distinct blueprint of proteolytic each organism uses. These digestion maps provide critical insight into *in vivo* protease cleavage patterns, determine where peptides are derived from within a proteome and allow differential digestion to be strains. In summary, DigestR was designed to address a key software gap in the study of protein catabolism.

## Acknowledgments

We thank Travis Bingeman for providing necessary support in developing the software tool. I.A.L. is supported by a UCalgary Research Excellence Chair and a grant from the Natural Sciences and Engineering Research Council of Canada (NSERC; DG04547). Proteomics data were acquired at the Calgary Metabolomics Research Facility, which is part of the Alberta Centre for Advanced Diagnostics (ACAD; PrairiesCan RIE #22734) and is supported by the International Microbiome Centre and the Canada Foundation for Innovation (CFI-JELF 34986).

## ASSOCIATED CONTENT

### Supporting Information

Documentation and code reference of the digestR R package can be found at https://www.lewisresearchgroup.org/software

### Author Contributions

The manuscript was written through contributions of all authors. All authors have given approval to the final version of the manuscript.

